# Dual regulation of the actin cytoskeleton by CARMIL-GAP

**DOI:** 10.1101/2021.03.09.434482

**Authors:** Goeh Jung, Miao Pan, Chris Alexander, Tian Jin, John A. Hammer

## Abstract

CARMIL (**Ca**pping protein **Ar**p2/3 **M**yosin **I** Linker) proteins are multi-domain scaffold proteins that regulate actin dynamics by regulating the activity of Capping Protein (CP). Here we characterize CARMIL-GAP, a *Dictyostelium* CARMIL isoform that contains a ~130 residue insert that, by homology, is a GTPase activating (GAP) domain for Rho-related GTPases. Consistently, this GAP domain binds *Dictyostelium* Rac1a and accelerates its rate of GTP hydrolysis. CARMIL-GAP concentrates with F-actin in phagocytic cups and at the leading edge of chemotaxing cells, and CARMIL-GAP null cells exhibit pronounced defects in phagocytosis and chemotactic streaming. Importantly, these defects are fully rescued by expressing GFP-tagged CARMIL-GAP in CARMIL-GAP null cells. Finally, rescue with versions of CARMIL-GAP that lack either GAP activity or the ability to regulate CP show that while both activities contribute significantly to CARMIL-GAP function, the GAP activity plays the bigger role. Together, our results add to the growing evidence that CARMIL proteins influence actin dynamics by regulating signaling molecules as well as CP, and that the continuous cycling of the Rho GTPase’s nucleotide state is often required to drive Rho-dependent biological processes.

**SUMMARY STATEMENT:** The assembly of actin filaments supports a wide array of fundamental cellular functions, including cell migration and phagocytosis. Actin assembly is controlled by a host of regulatory proteins, with Capping Protein being one of the most important. Capping Protein is in turn regulated by the CARMIL family of proteins. Actin assembly is also controlled by signaling pathways that often converge on Rho-related GTPases like Rac1. These GTPases cycle between an active, GTP-bound state and an inactive, GDP-bound state. Guanine nucleotide exchange factors (GEFs) and guanine nucleotide activating proteins (GAPs) drive Rho-related GTPases to their GTP-bound and GDP-bound states, respectively. Here we characterized a version of CARMIL that contains within it a GAP domain for Rac1. We show that CARMIL-GAP supports the actin-based processes of cell migration and phagocytosis. We also show that while CARMIL-GAP’s ability to regulate Capping Protein and the nucleotide state of Rac1 are both important for its cellular functions, its ability to regulate Rac1 via its GAP domain plays the bigger role. Finally, our data support the emerging concept that the continuous cycling of Rho GTPases between their GTP-bound and GDP-bound states is often required to drive Rho-dependent biological processes.

## INTRODUCTION

*Dictyostelium* CARMIL and its ortholog Acan 125 in *Acanthamoeba* are the founding members of a class of proteins that regulate Capping Protein (CP), the primary actin filament barbed end capping protein in most if not all Eukaryotic cells and a central player in the assembly, organization and dynamics of the actin cytoskeleton (Edwards et al., 2014; Jung et al., 2001; Xu et al., 1995). Identified based on their interaction with the SH3 domains of type 1 myosins (Jung et al., 2001; Xu et al., 1995), these ~1100-residue, multidomain scaffold proteins and their metazoan counterparts contain a ~50 residue domain that binds CP with nanomolar affinity (Remmert et al., 2004; Uruno et al., 2006; Yang et al., 2005). This domain, referred to in the literature as either CARMIL Homology Domain 3 (CAH3) or Capping Protein Interacting Domain (CPI), exerts two dramatic and interrelated biochemical effects on CP. First, when bound to CP, CPI reduces CP’s affinity for the fast-growing barbed end of the actin filament from 0.1 nM to ~30 nM (Uruno et al., 2006; Yang et al., 2005). This reduction in affinity equates to a reduction in CPs’s half-life on the barbed end from ~30 minutes to ~10 seconds. Second, when added to CP-capped actin filaments, CPI dramatically accelerates the dissociation of CP from the barbed end such that, at saturation, CPI reduces CP’s half-life on the barbed end from ~30 minutes to ~10 seconds (Fujiwara et al., 2010). Importantly, these two biochemical effects are variations of the same underlying mechanism, which is that the binding of CPI to CP alters CP’s conformation in such a way as to reduce its affinity for the barbed end by several hundred fold (Hernandez-Valladares et al., 2010; Kim et al., 2012; Takeda et al., 2010; Zwolak et al., 2010b). Stated another way, CPI’s allosteric effect on CP serves to both reduce the affinity of preformed CP:CPI complexes for the barbed end, and promote uncapping when CPI is added to filaments already capped with CP.

While the biochemical effects exerted by CPI on CP *in vitro* would suggest that CARMIL proteins function as CP antagonists *in vivo,* the truth may well be the opposite. To understand how this is possible, one must consider CARMIL function in cells in the context of a second direct regulator of CP known as V-1. This ubiquitously-expressed, small ankyrin repeat protein binds CP 1:1 with ~20 nM affinity to prevent it from capping the barbed end (Bhattacharya et al., 2006; Takeda et al., 2010; Zwolak et al., 2010a). Importantly, V-1 is present in the cytoplasm at a 3- to 4-fold molar excess over CP (Fujiwara et al., 2014; Jung et al., 2016). Given this, and given V-1’s affinity for CP, one would predict that ~99% of cellular CP would be sequestered by V-1, barring regulation. Critically, CARMIL’s CPI domain provides a powerful counter to V-1’s sequestering activity by robustly catalyzing an exchange reaction that converts CP:V-1 complexes into CP:CPI complexes (Fujiwara et al., 2014; Johnson et al., 2018; Takeda et al., 2010). In this way, CARMIL proteins convert inactive CP (CP:V-1) into a version (CP:CPI) with moderate (i.e. ~30 nM) affinity for the barbed end. In other words, in the face of pervasive sequestration of CP by V-1, CARMIL proteins should serve as activators rather than inhibitors of CP. These findings, when combined with other data regarding the localization and activation of CARMIL at the plasma membrane (Fujiwara et al., 2014; Zwolak et al., 2013), argue that CARMIL and V-1 cooperate at the leading edge of cells to promote Arp2/3 complex-dependent branched actin network assembly there by promoting weak barbed-end capping(Fujiwara et al., 2014). Consistent with this model, estimates of CP’s half-life on barbed ends near the plasma membrane *in vivo* (~2 to 15 seconds) (Iwasa and Mullins, 2007; Miyoshi et al., 2006) are much closer to the half-life of the CP:CPI complex on the barbed end (~10 seconds) than to the half-life of CP alone (~30 minutes) (Fujiwara et al., 2010). This model is also consistent with evidence that CARMIL proteins promote lamellipodia formation (Edwards et al., 2015; Jung et al., 2001; Liang et al., 2013), that cells forced to express a version of CP that can cap barbed ends but cannot see the CPI motif exhibit a CP-knockdown phenotype (Edwards et al., 2015), and that the defects in actin organization and dynamics exhibited by cells devoid of V-1 or over-expressing V-1 both demonstrate that V-1 regulates CP activity *in vivo* (Jung et al., 2016).

While this brief overview of CARMIL’s CP1 domain emphasizes the role played by CARMIL proteins in regulating actin assembly by regulating CP activity, there is growing evidence that these scaffold proteins also regulate actin assembly by regulating signaling pathways (reviewed in (Stark et al., 2017)). One clear example is the *C.elegans* CARMIL homologue CRML-1, which negatively regulates neuronal growth cone migration by binding to and inhibiting UNC-73, the *C.elegans* homologue of Trio, a guanine nucleotide exchange factor (GEF) for Rac and Rho (Vanderzalm et al., 2009). Consistent with this finding, immunoprecipitates of CARMIL-1 from human fibroblasts contain Trio (Liang et al., 2009), and the CARMIL-2 interactome in T cells contains two GEFs for Rho-related GTPases (VAVl and DOCKS) (Liang et al., 2013). Interestingly, CARMIL-2 also serves to link the cell surface receptor CD28 in T cells to the adaptor molecule CARMA1, which then collaborates with PKCθ to promote full T cell activation by activating the transcription factor NF-KB (Liang et al., 2013; Roncagalli et al., 2016).

In our 2001 study of *Dicytostelium* CARMIL we showed that it binds myosin 1, CP and the Arp2/3 complex, and that it concentrates with them in several actin-rich structures, most notably macropinocytic projections on the dorsal surface of vegetative cells and pseudopods at the leading edge of starved, aggregating cells (Jung et al., 2001). Consistent with these localizations, CARMIL null cells exhibited pronounced defects in macropinocytosis and chemotactic aggregation (Jung et al., 2001). While the CARMIL isoform examined in that study was considered at the time to be the only CARMIL isoform in *Dictyostelium,* the subsequent completion of the *Dictyostelium* genome sequence revealed the presence of a second CARMIL gene. We call this second isoform CARMIL-GAP because the protein contains an apparent guanine nucleotide activating (GAP) domain for Rho-related GTPases in addition to all the normal domains present in CARMILs. Here we characterized the localization and cellular functions of CARMIL-GAP, and we used complementation to parse out the relative contributions made by its CPI and GAP domains to its cellular functions. Together, our results add to the growing evidence that CARMIL proteins regulate actin dynamics by regulating signaling molecules as well as CP, although in the case of CARMIL-GAP this dual regulation is accomplished not in trans but by this one protein. Finally, our results support the emerging concept (Denk-Lobnig and Martin, 2019) that the continuous cycling of Rho GTPases between their GTP and GDP bound states through the coordinated action of their GEFs and GAPs is often required to drive Rho-dependent biological processes forward.

## RESULTS

### CARMIL-GAP contains functional GAP and CPI domains

Alignment of the amino acid sequence of *Dictyostelium* CARMIL-GAP with that of *Dictyostelium* CARMIL (Figure S1) shows that CARMIL-GAP contains, in addition to the five domains found in all CARMIL proteins (a pleckstrin homology-like (PH-like) domain, a leucine rich repeat (LRR) domain, a homo-dimerizing (HD) domain, a proline-rich (Pro) domain, and a CPI domain (Edwards et al., 2013)), a ~145-residue sequence inserted between its HD and Pro domains (Figure 1, Panel A). Blast searches revealed that this insert is homologous to the GAP domains present in GTPase activating proteins for Rho-related GTPases such as Cdc42-GAP and ARHGAP4 (Barfod et al., 1993; Vogt et al., 2007) (Figure 1B, Panel B). Importantly, this putative GAP domain in CARML-GAP contains an arginine residue present in all GAP domains that is required for robust GAP activity (residue 737; highlighted red in Figure 1, Panel B), as well as polar residues at positions 780, 841 and 849 (highlighted blue in Figure 1, Panel B) that are required for stabilization of the Rho-GTPase’s switch domain during GAP-stimulated GTP hydrolysis (Nassar et al., 1998; Rittinger et al., 1997; Scheffzek et al., 1997). Together, these sequence elements suggest that CARMIL-GAP functions at least in part as a GAP for a Rho-related GTPase.

**Figure 1.**
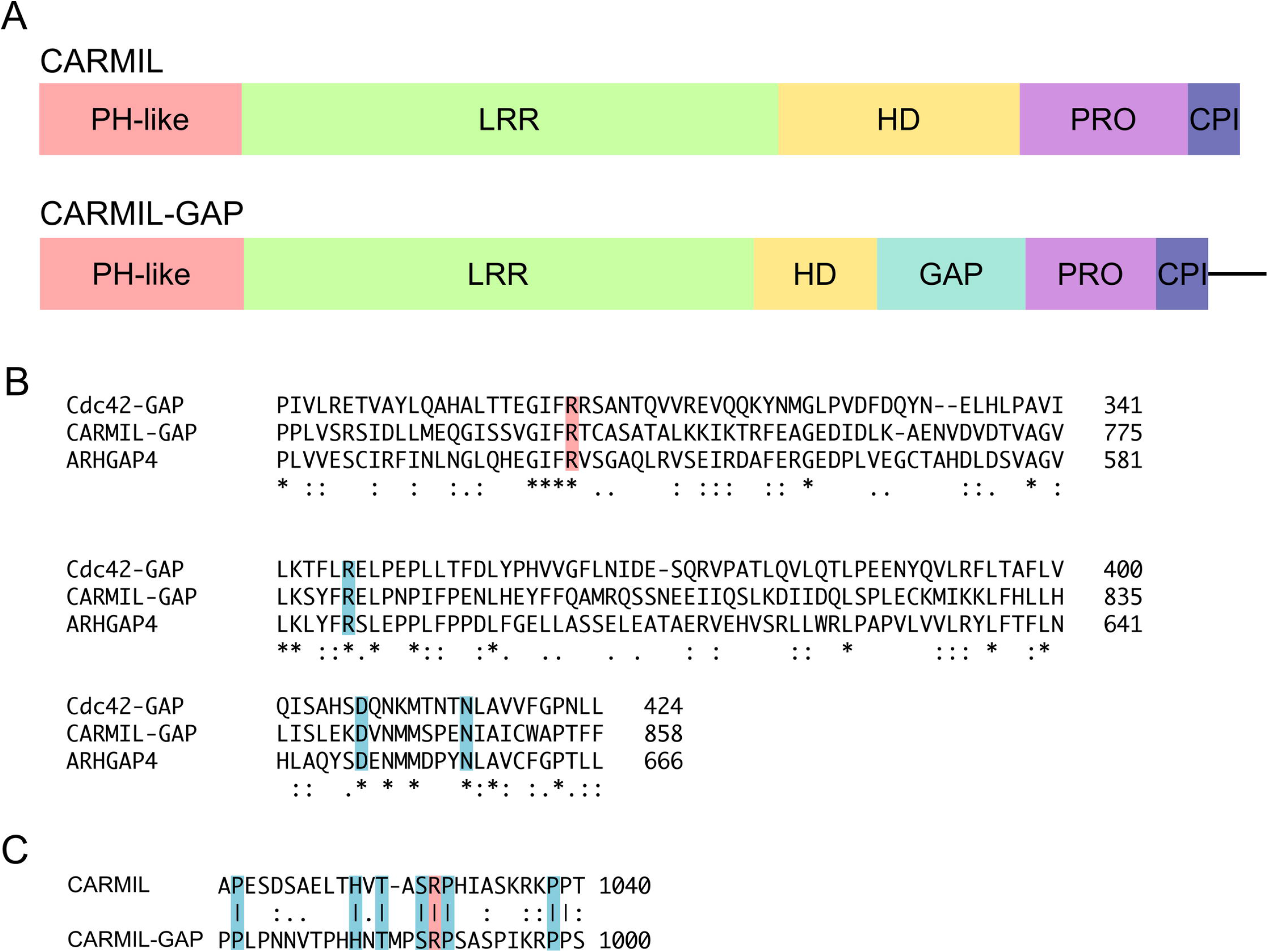
Domain organization of CARMIL-GAP. (A) Cartoon depicting the domain organization of CARMIL-GAP versus CARMIL (Jung et al., 2001) (PH-like, pleckstrin homology-like domain (Zwolak et al., 2013); LRR, leucine-rich repeat domain; HD, homodimerizing domain (Zwolak et al., 2013); Pro, proline-rich domain; CPI, Capping Protein Interaction domain). (B) Alignment of the putative GAP domain of CARMIL-GAP with the GAP domains of human Cdc42-GAP (Vogt et al., 2007) and human ARHGAP4 (Barfod et al., 1993). The conserved arginine at position 737 in CARMIL-GAP, 305 in Cdc42-GAP, and 544 in ARHGAP4 (shaded red) is required for robust GAP activity. The conserved residues shaded blue are required for stabilization of the Rho-GTPase’s switch domain during GAP-stimulated GTP hydrolysis. (C) Alignment of the CPI domains in CARMIL and CARMIL-GAP. The arginine at position 1029 in CARMIL and 989 in CARMIL-GAP (shaded red) is essential for binding CP with high affinity (Edwards et al., 2014; Uruno et al., 2006; Yang et al., 2005). The residues shaded blue were identified by site directed mutagenesis of *Acanthamoeba* CARMIL’s CPI domain as contributing significantly to its anti-CP activity (Uruno et al., 2006). Lines indicate identity, colons indicate highly conservative substitutions, and dots indicate moderately conservative substitutions.

To seek evidence that the putative GAP domain in CARMIL-GAP functions as a GAP, we first sought to identify the Rho-related GTPase(s) that interacts with it. In terms of candidate Rho-related GTPases, *Dictyostelium* possesses six Rac GTPases (Rac1a, Rac1b, Rac1c, RacB, RacF1, and RacF2), thirteen Rac-like GTPases (RacC, RacD, RacE, RacG, RacH, Rac1, RacJ, Rac1, RacM, RacN, RacO, RacP, and RacQ), and one RhoBTB homolog (RacA), but no obvious Rho or Cdc42 family GTPases (Rivero and Xiong, 2016). To identify the Rac isoform(s) that interacts with CARMIL-GAP, its isolated GAP domain (specifically residues 715 to 858) was expressed as a GST fusion, bound to Glutathione Sepharose 4B, and incubated with *Dictyostelium* cell lysates. After extensive washes, the bound material was eluted with high salt buffer, concentrated, digested with trypsin, and the digest subjected to mass spectrometry/sequence analysis. In terms of Rho-related GTPases, the curated list of bound proteins (Table S1; see the legend for details) contained only two Rho-related GTPases: the Rac-like GTPase RacE (10 distinct peptides) and Rac1 (6 distinct peptides; Figure 2, Panel A). With regard to Rac1, while peptides 1, 3 and 4 are present in all three Rac1 isoforms (Rac1a, Rac1b, and Rac1c), and peptide 5 is present in both Rac1a and Rac1b, peptides 2 and 6 are present only in Rac1a. These two Rac1a-specific peptides also comprised 6 of the 13 Rac1-related peptides present in the non-curated list of bound proteins. The most straightforward interpretation of these results is that CARMIL-GAP’s GAP domain interacts preferentially if not exclusively with the 1a isoform of *Dictyostelium* Rac1 (although see Discussion). Given the phenotype of CARML-GAP null cells (see below), which do not exhibit a defect in cytokinesis when grown in suspension (a process regulated by RacE; reviewed in Rivero and Xiong, 2016)), but do exhibit a pronounced defect in phagocytosis (a process regulated by Rac1a; reviewed in Rivero and Xiong, 2016)), we focused on the role of CARMIL-GAP’s GAP domain in regulating the nucleotide state of Rac1a.

**Figure 2.**
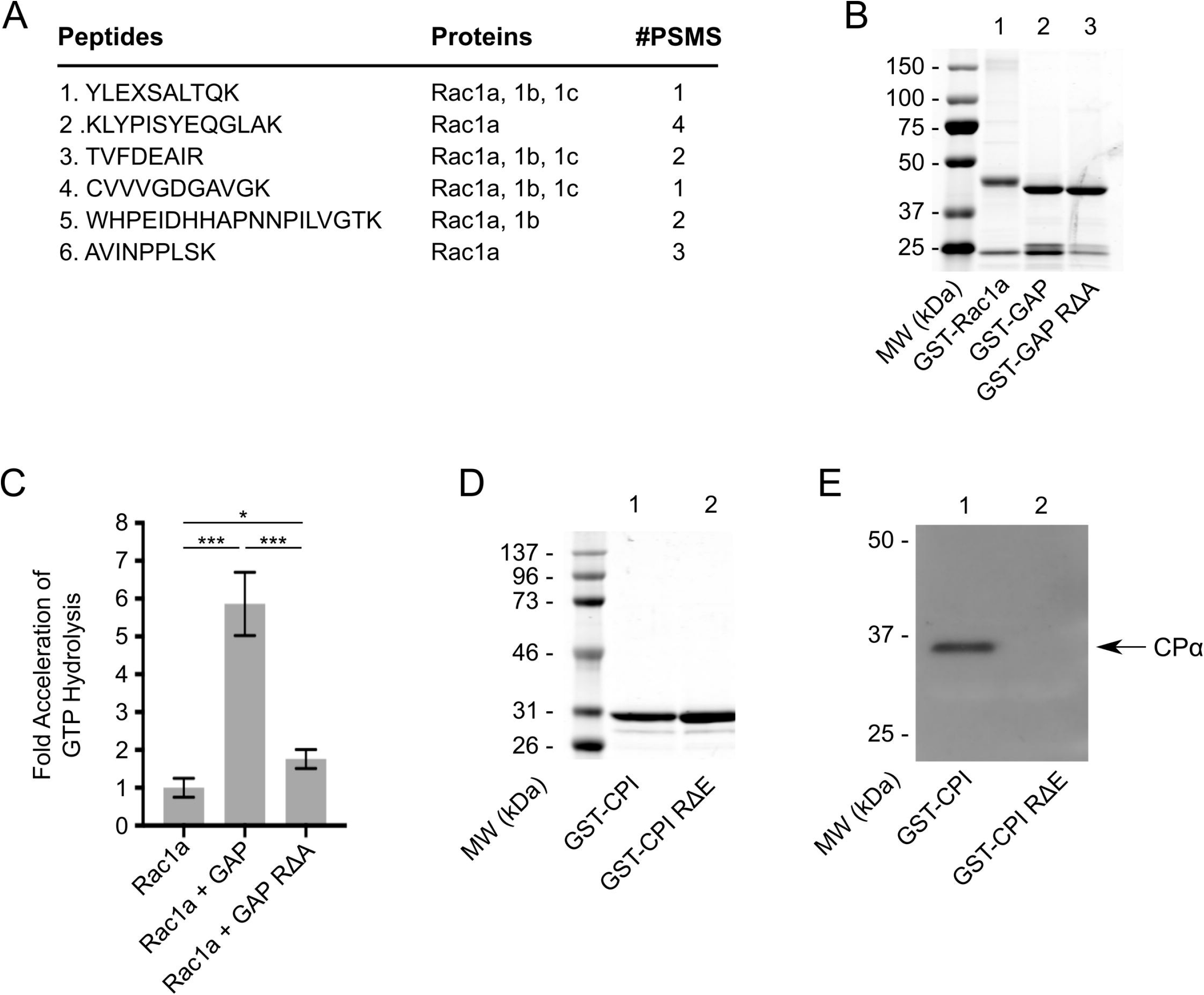
CARMIL-GAP contains functional GAP and CPI domains. (A) Shown are the six unique Rac1 peptides obtained by mass spec analysis of the GST-GAP domain pull down, the Rac1 isoform they were found in, and the frequency with which each was identified (PSMs). (B) Coomassie Blue-stained gel of purified GST-Rac1a (lane 1), GST-GAP (lane 2), and GST-GAP-RΔA (lane 3). (C) Shown is the fold activation of GST-Rac1a’s GTP hydrolysis rate (mean normalized to 1.0) by GST-GAP or GST-GAP-RΔA (Rac1a, 1.00 ± 0.25; Rac1a + GAP, 5.80 ± 0.84; Rac1a + GAP RΔA, 1.74 ± 0.25; N=3). (D) Coomassie Blue-stained gel of purified GST-CPI and GST-CPI-RΔA. (E) Western blot of the material eluted from GST-CP1 beads (lane1) and GST-CPI-RΔA beads (lane 2) after incubation with whole cell extracts, and probed with an antibody to the alpha subunit of *Dictyostelium* Capping Protein.

To obtain direct evidence that the GAP domain in CARMIL-GAP functions to accelerate the rate of GTP hydrolysis by Rac1a, we expressed as N-terminally tagged GST fusion proteins Rac1a (GST-Rac1a), the GAP domain (GST-GAP), and a version of the GAP domain in which the conserved arginine residue required for robust GAP activity in known GAPs was changed to an alanine residue (GST-GAP-RΔA). Figure 2, Panel B, shows these three fusion proteins following purification. Of note, specific conditions were used to obtain GST-Rac1a in its GTP-bound form prior to performing the GAP assay (see Methods). To measure the effect of CARMIL-GAP’s GAP domain on the rate of GTP hydrolysis by Rac1a, 10 μM of GST-Rac1a was incubated for 10 minutes at 20°C with either 2.5 μM GST-GAP or GST-GAP-RΔA, at which point the amount of free phosphate in solution (a measure of GTP hydrolysis by GST-Rac1a) was determined by measuring the absorbance of the dye CytoPhos (see Methods). Figure 2, Panel C, shows that the addition of GST-GAP increased Rac1a’s intrinsic rate of GTP hydrolysis (normalized to a value of 1.0) by an average of 5.8 fold. In contrast, the addition of GST-GAP-RΔA increased Rac1a’s rate of GTP hydrolysis by only 1.7 fold. Together, these data argue that the GAP domain in CARML-GAP is functional, and that CARMIL-GAP likely functions as a GAP for Rac1a in *Dictyostelium.*

As discussed in the Introduction, CARMIL proteins are best known for their interaction with and regulation of CP, which is mediated by their CPI domain. By sequence alignment (Figure 1, Panel C), CARMIL-GAP possesses a typical CPI domain containing the invariant arginine that is essential for CP interaction (residue 989; highlighted red in Figure 1, Panel C), as well as other residues that potentiate CP binding and regulation (highlighted blue in Figure 1, Panel C) (McConnell et al., 2020; Uruno et al., 2006). To demonstrate that CARMIL-GAP does indeed bind CP, residues spanning its CPI domain (residues 963-1005) were expressed as an N-terminally tagged GST fusion protein (GST-CPI), bound to Glutathione Sepharose 4B resin, incubated with *Dictyostelium* cell lysates and, after extensive washes, the bound proteins were eluted with high salt buffer. As a negative control, a parallel binding reaction was performed using a version of this CPI fusion protein in which the essential arginine at position 989 was changed to a glutamate (GST-CPI-RΔE). Figure 2, Panel D, shows these two fusion proteins following purification. A Western blot of the bound proteins probed with an antibody to the alpha subunit of *Dictyostelium* CP showed that CP was present in the material eluted from GST-CPI (Figure 2, Panel E, lane 1) but not in the material eluted from GST-CPI-RΔE (Figure 2, Panel E, lane 2). These results argue that CARMIL-GAP also functions as a regulator of CP.

### CARMIL-GAP null cells grow normally in liquid media and do not exhibit defects in macropinocytosis and cell division

Western blots of *Dictyostelium* whole cell extracts probed with an antibody raised against the C-terminal 19 residues of CARMIL-GAP show that it is expressed in both vegetative cells and starved, developing cells (Figure S2, Panel A). This result, together with the biochemical evidence above that it likely regulates CP, a central player in actin assembly, and Rac1, a central player in the regulation of actin assembly, argues that CARMIL-GAP may regulate actin-dependent processes occurring in both of these physiological states (e.g. macropinocytosis, cell division and phagocytosis in vegetative cells, and cell motility in starved, aggregating cells). To define CARMIL-GAP’s physiological significance in both vegetative and starved cells, and to permit dissection of its *in vivo* functions by complementation, we created CARMIL-GAP null cells using homologous recombination. Briefly, AX3 cells were transformed with a linear gene disruption fragment containing the selectable marker blasticidin S flanked by portions of the CARMIL-GAP gene, and single, blasticidin S-resistant cells were cloned by serial dilution (see Methods for details). Figure 3, Panel A, shows a Western blot of whole cell extracts prepared from wild type (WT) cells and the two independent CARMIL-GAP knockout (KO) cell lines (M1 and M2) that we used interchangeably in this study.

**Figure 3.**
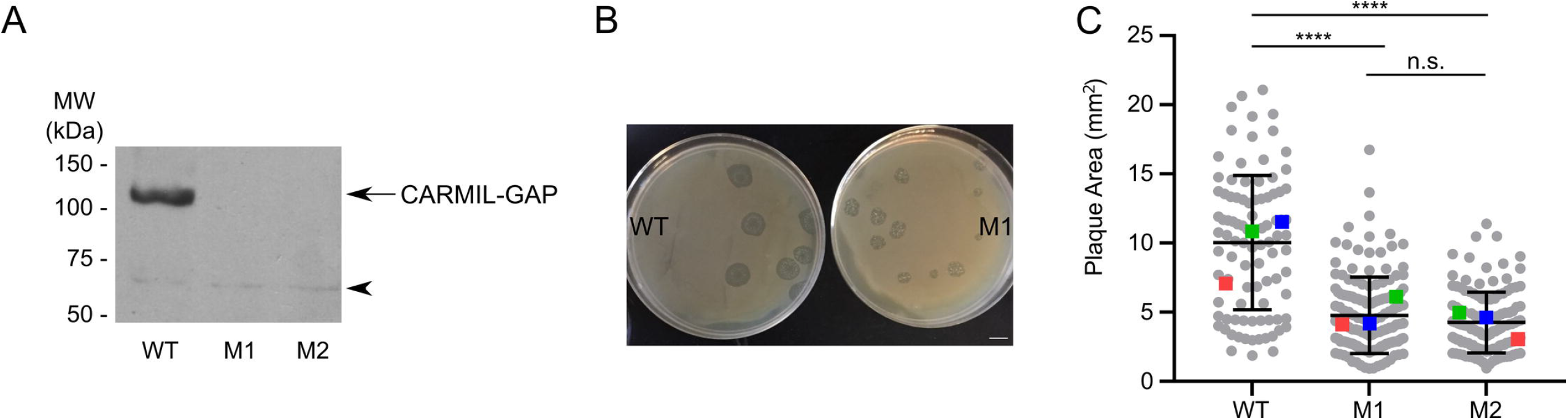
CARMIL-GAP null cells grown on bacterial lawns make significantly smaller plaques. (A) Western blot of whole cell extracts prepared from equal numbers of control AX3 cells (WT) and CARMIL-GAP null cell lines M1 and M2 probed with an antibody to CARMIL-GAP. The cross reacting band at ~63 kDa (see arrowhead) serves as a loading control (see also Figure S3, lanes 1 and 2, which included an actin loading control). (B) Representative examples of WT and M1 KO cells grown on a lawn of **K.* aerogenes* for five days (plaque assay). (Cl Quantitation of plaque size at day five for WT, M1 KO, and M2 KO cells (see also Table 1). The number of plaques scored over three independent experiments is shown at the bottom of each bar, and the mean values for the three experiments performed are indicated by the red, green and blue squares.

In terms of baseline data in vegetative cells, CARMIL-GAP KO cells grew at normal rates in HL5 liquid media (Figure S2, Panel B). Consistently, CARMIL-GAP KO cells exhibited normal rates of macropinocytosis, the mechanism by which axenic strains gain nutrients when grown in liquid media (Figure S2, Panel C) (Hacker et al., 1997). Moreover, vegetative CARMIL-GAP KO cells do not exhibit a reduction in steady state F-actin content (Figure S2, Panel D). Finally, vegetative CARMIL-GAP KO cells do not exhibit a significant defect in cytokinesis based on measuring the fraction of cells grown in suspension that contain more than one nuclei (Figure S2, Panel E). Taken together, these baseline data indicate that CARMIL-GAP does not play a significant role in either macropinocytosis or cell division, two major actin-dependent processes occurring in vegetative cells.

**Table 1.**
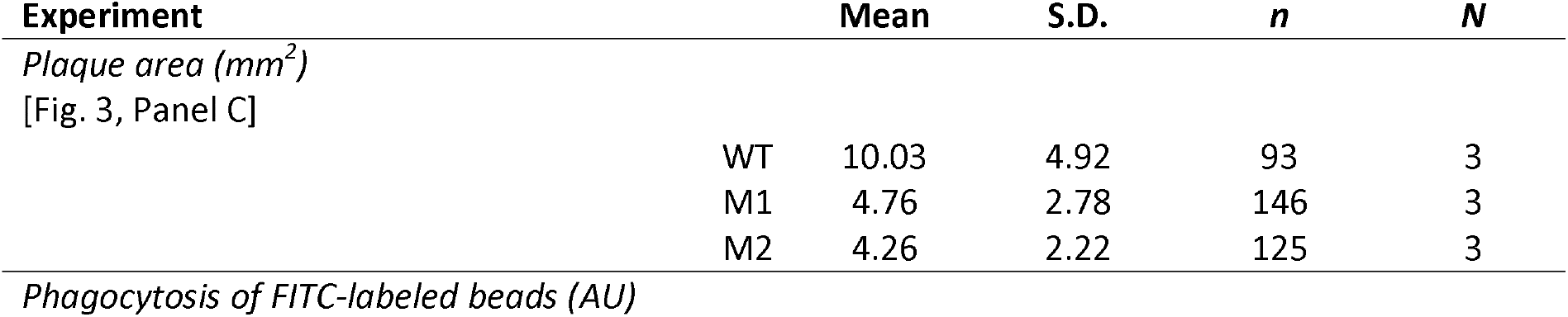
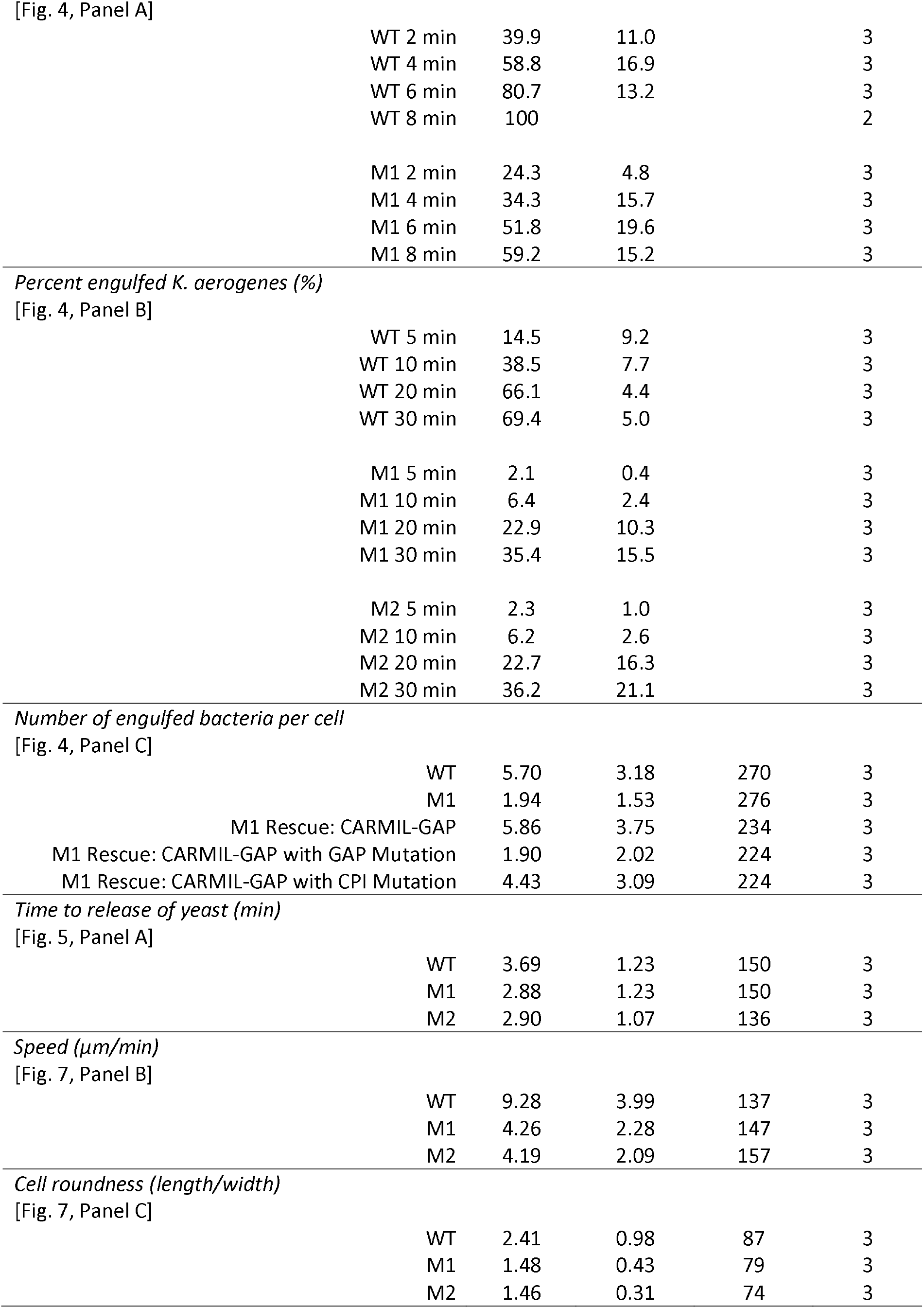
Means, standard deviations and N values.

| Experiment |  | Mean | S.D. | n | N |
| --- | --- | --- | --- | --- | --- |
| *Plaque area (mm<sup>2</sup>)* |  |  |  |  |  |
| [Fig. 3, Panel C] |  |  |  |  |  |
|  | WT | 10.03 | 4.92 | 93 | 3 |
|  | M1 | 4.76 | 2.78 | 146 | 3 |
|  | M2 | 4.26 | 2.22 | 125 | 3 |
| *Phagocytosis of FITC-labeled beads (AU)* |  |  |  |  |  |

|  |  |  |  |
| --- | --- | --- | --- |
| WT 2 min | 39.9 | 11.0 | 3 |
| WT 4 min | 58.8 | 16.9 | 3 |
| WT 6 min | 80.7 | 13.2 | 3 |
| WT 8 min | 100 |  | 2 |
| M1 2 min | 24.3 | 4.8 | 3 |
| M1 4 min | 34.3 | 15.7 | 3 |
| M1 6 min | 51.8 | 19.6 | 3 |
| M1 8 min | 59.2 | 15.2 | 3 |

|  |  |  |  |
| --- | --- | --- | --- |
| WT 5 min | 14.5 | 9.2 | 3 |
| WT 10 min | 38.5 | 7.7 | 3 |
| WT 20 min | 66.1 | 4.4 | 3 |
| WT 30 min | 69.4 | 5.0 | 3 |
| M1 5 min | 2.1 | 0.4 | 3 |
| M1 10 min | 6.4 | 2.4 | 3 |
| M1 20 min | 22.9 | 10.3 | 3 |
| M1 30 min | 35.4 | 15.5 | 3 |
| M2 5 min | 2.3 | 1.0 | 3 |
| M2 10 min | 6.2 | 2.6 | 3 |
| M2 20 min | 22.7 | 16.3 | 3 |
| M2 30 min | 36.2 | 21.1 | 3 |

|  |  |  |  |  |
| --- | --- | --- | --- | --- |
| WT | 5.70 | 3.18 | 270 | 3 |
| M1 | 1.94 | 1.53 | 276 | 3 |
| M1 Rescue: CARMIL-GAP | 5.86 | 3.75 | 234 | 3 |
| M1 Rescue: CARMIL-GAP with GAP Mutation | 1.90 | 2.02 | 224 | 3 |
| M1 Rescue: CARMIL-GAP with CPI Mutation | 4.43 | 3.09 | 224 | 3 |

|  |  |  |  |  |
| --- | --- | --- | --- | --- |
| WT | 3.69 | 1.23 | 150 | 3 |
| M1 | 2.88 | 1.23 | 150 | 3 |
| M2 | 2.90 | 1.07 | 136 | 3 |

|  |  |  |  |  |
| --- | --- | --- | --- | --- |
| WT | 9.28 | 3.99 | 137 | 3 |
| M1 | 4.26 | 2.28 | 147 | 3 |
| M2 | 4.19 | 2.09 | 157 | 3 |

|  |  |  |  |  |
| --- | --- | --- | --- | --- |
| WT | 2.41 | 0.98 | 87 | 3 |
| M1 | 1.48 | 0.43 | 79 | 3 |
| M2 | 1.46 | 0.31 | 74 | 3 |

### CARMIL-GAP null cells grown on bacterial lawns make significantly smaller plaques, suggesting a defect in phagocytosis

To explore the possibility that CARMIL-GAP plays a significant role in phagocytosis, another major actin-dependent process exhibited by vegetative cells, we initially measured the rate at which cells create bacteria-free plaques when grown in the presence of an even lawn of bacteria as the nutrient source. To accomplish this, WT and CARMIL-GAP KO lines M1 and M2 where seeded at low density with living *Klebsie/Ja aerogenes* bacteria on agar plates made using five-fold diluted HL5. *Dictyostelium* will not grow on such agar plates alone, so their ability to grow in the presence of the bacteria, which is scored as the size of the plaques that form after five days, should be due to their ability to phagocytose the bacteria. Figure 3, Panel B, shows that KO line M1 made significantly smaller plaques than WT cells. Consistently, quantitation showed that both KO lines produced plaques that were about 35% the size of plaques produced by WT cells (Figure 3, Panel C; see also Table 1). These results suggest that CARMIL-GAP plays a significant role in supporting the actin-dependent process of phagocytosis.

### CARMIL-GAP null cells exhibit a significant defect in phagocytosis

The smaller plaque size exhibited by CARMIL-GAP null cells grown on bacterial lawns could result from defects in processes other than phagocytosis (e.g. a defect in the ability to digest the bacteria; (Buckley et al., 2016; Pan et al., 2016). Given this, we sought more direct measures of phagocytic ability. As a first attempt, we measured the initial rate of uptake of fluorescent 1 μm polystyrene beads as the phagocytic substrate. Figure 4, Panel A, shows that CARMIL-GAP KO line M1 exhibited about a 40% decrease in bead uptake over 8 minutes relative to WT cells (see also Table 1). To provide a more physiological measure of phagocytosis, we used a FACS-based assay (Pan et al., 2018b; Pan et al., 2016) that measures the phagocytosis of pHrodo Red dye-labeled *Klebsiella aerogenes,* whose fluorescence within phagosomes increases dramatically upon the acidification of the phagosome. Figure 4, Panel B, shows that the percentage of M1 and M2 cells in suspension that had taken up bacteria over a 30 minute incubation was less than half that of WT cells (see also Table 1). Moreover, quantitative confocal imaging (Pan et al., 2018b; Pan et al., 2016) showed that adherent M1 cells internalized ~67% fewer bacteria than WT cells after a 15 minute incubation (Figure 4, Panel C, and Table 1; compare “M1” to “WT”; see also Panel D for representative images of these two samples). Importantly, the re-expression of CARMIL-GAP (as a GFP fusion) in M1 cells fully rescued this defect in phagocytosis (Figure 4, Panel C and Table 1; compare rescue with “CARMIL-GAP” to “WT”; see also Panel D for representative images of these two samples, Figure S3, lane 3, which shows that these cells make a protein corresponding in size to GFP-CARMIL-GAP, and Figure S4, Panels A1-A4, which shows that GFP-tagged CARMIL-GAP localizes to phagocytic cups in complemented null cells). Together, these results argue that the defect in phagocytosis is indeed caused by the loss of CARMIL-GAP.

**Figure 4.**
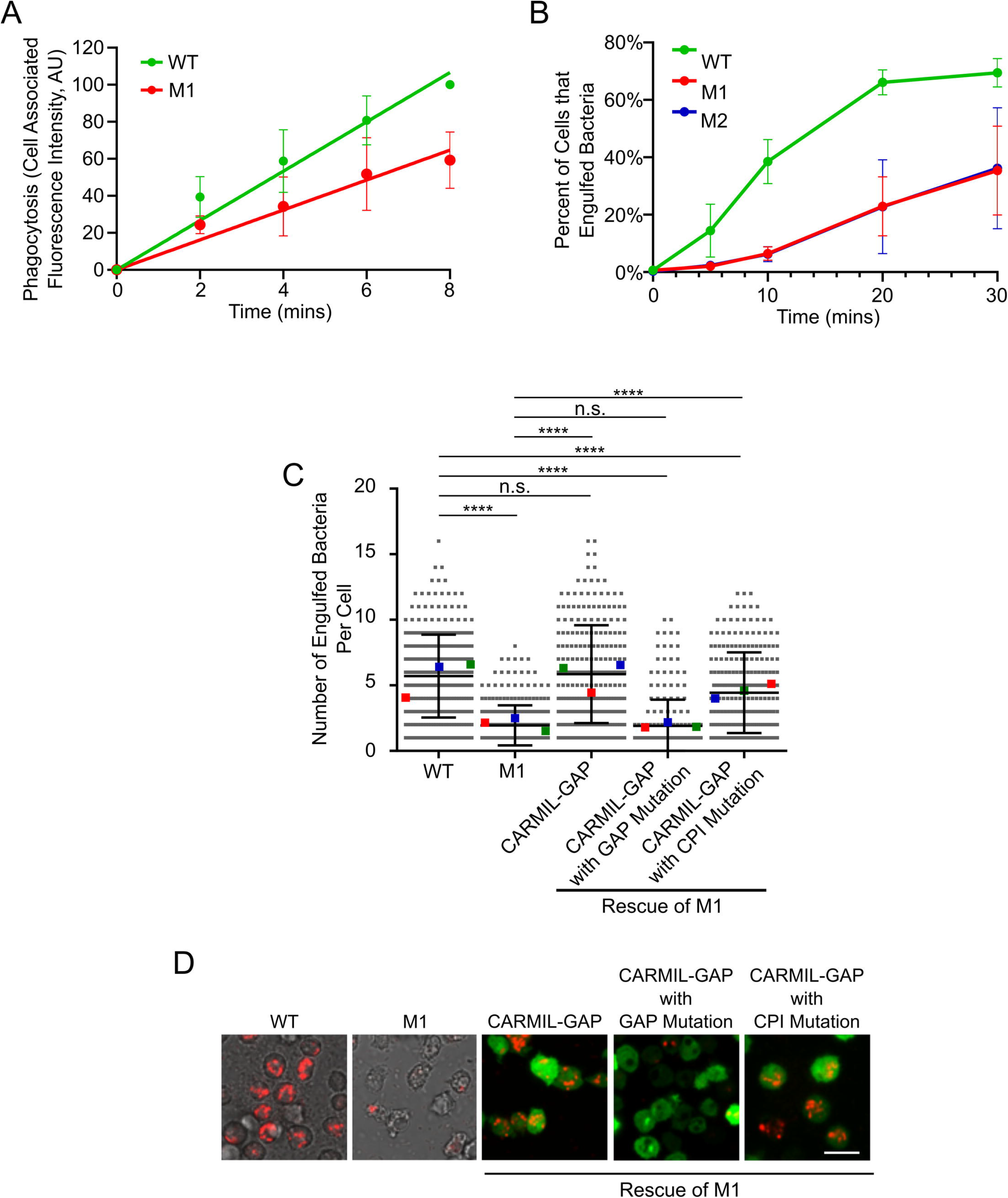
CARMIL-GAP null cells exhibit a significant defect in the phagocytosis of bacteria. (A) Shown is the initial rate of uptake by phagocytosis of 1 μm FITC-labeled polystyrene beads by WT and M1 KO cells, presented as cell-associated fluorescence (in arbitrary units; AU) (see also Table 1). (B) Shown are the percentages of WT, M1 KO, and M2 KO cells that had engulfed pHrodo Red dye-labeled *K. aerogenes* by 5, 10, 20 and 30 minutes, as determined by FACs (see also Table 1). (C) Shown are the numbers of *K. aerogenes* engulfed per cell after 15 minutes of incubation, as determined by quantitative confocal microscopy, for WT and M1 KO cells, and for M1 KO cells that were complemented with GFP-tagged versions of CARMIL-GAP, CARMIL-GAP containing the GAP domain mutation, or CARMIL-GAP containing the CPI domain mutation (see also Table 1). The mean values for the three experiments performed are indicated by the red, green and blue squares. (D) Shown are representative images of the five cell samples quantified in Panel D (red, bacteria; green, GFP-CARMIL). Mag bar: 10 μm.

To supplement these results, we used a confocal microscopy-based assay to follow the phagocytosis of fluorescently-labeled yeast, which represent a significantly larger phagocytic substrate than bacteria. Imaging was initiated upon contact with a yeast particle and continued every 5 seconds for 30 minutes after contact. For both WT cells and KOs M1 and M2, most of these contacts failed to lead to internalization, with WT, M1 and M2 cells releasing bound yeast particles on average 3.7 ± 1.2, 2.8 ± 1.2, and 2.9 ± 1.1 minutes after contact, respectively (Figure 5, Panel A; see also Table 1). Figure 5, Panels C1-C4, show a representative example of such a failed phagocytic event in a WT cell (see also Movie 1). Despite the high failure rate, a subset of WT and KO cells possessed a fluorescent yeast particle 15 minutes after initial contact, or roughly five times longer than the average time for yeast particle release. Specifically, 26.3% of WT cells (54 out of 205 cells), 10.7% of M1 KO cells (18 out of 168 cells), and 11.3% for M2 KO cells (17 out of 150 cells) still possessed a fluorescent yeast particle 15 minutes after initial contact (Figure 5, Panel B). Figure 5, Panels D1-D4, show a representative example of a successful phagocytic event in a M1 KO cell (see also Movie 2). Further evidence that (see Methods for additional details). We conclude, therefore, that the loss of CARMIL-GAP results in a ~58% decrease in the efficiency of yeast phagocytosis. Finally, staining of WT cells in the process of phagocytosing an unlabeled yeast particle showed that CARMIL-GAP accumulates in the phagocytic cup along with F-actin (Figure 5, Panels E1-E4). Together, the above results indicate that CARMIL-GAP plays a major role in the actin-dependent process of phagocytosis.

**Figure 5.**
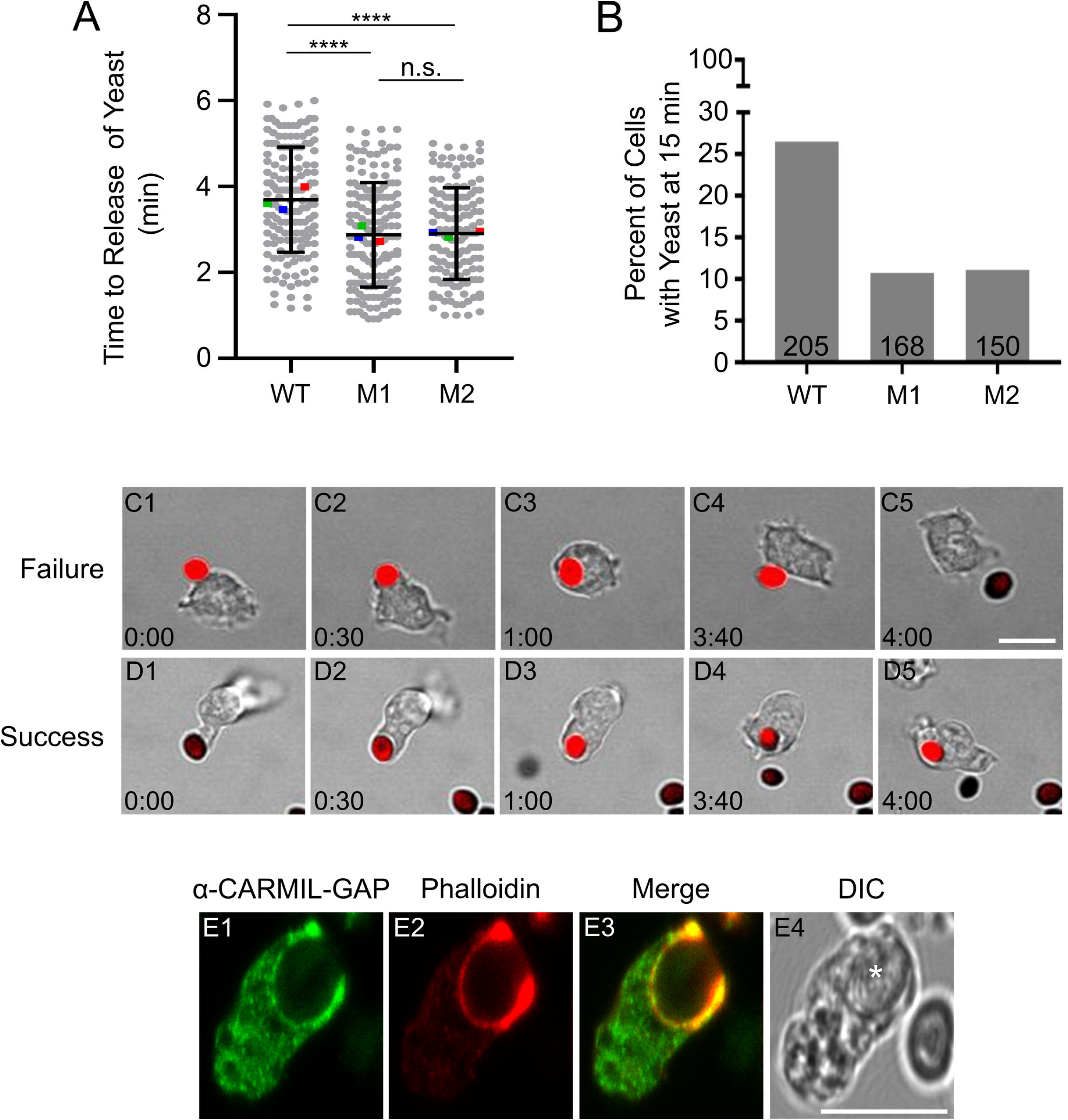
CARMIL-GAP null cells exhibit a significant defect in the phagocytosis of yeast. (A) Shown are times in minutes to release of bound yeast particles for WT, M1 KO and M2 KO cells (see also Table 1). The mean values for the three experiments performed are indicated by the red, green and blue squares. (B) Percent of WT, M1 KO and M2 KO cells that still retained a yeast particle 15 minutes after initial contact. The total number of cells scored from three independent experiments, is indicated at the bottom of each bar. (Cl-C4) Still images taken from a representative movie of a WT cell where a bound yeast particle was released. (Dl-04) Still images taken from a representative movie of an M1 KO cell where a bound yeast particle was successfully internalized. (El-E4) Image of a representative *Dictyostelium* cell engulfing an unlabeled yeast particle (white asterisk in the DIC image in E4) and stained for endogenous CARMIL-GAP (El) and F-actin (E2) (E3 shows the merged image). Mag bar: 10 μm.

### CARMIL-GAP null cells exhibit a defect in chemotactic streaming

Given that CARMIL-GAP is expressed in starved cells as well as vegetative cells, we next asked if it plays a significant role in the actin-dependent process of chemotactic aggregation that is initiated by starvation. Specifically, starvation sets in motion a developmental program that drives the coalescence of ~100,000 cells to form a stalk with a spore-filled head. Cell coalescence is driven by the migration of cells towards an extracellular gradient of cAMP that is generated initially by a small number of pioneer cells. Individual amoeba initially chemotaxis towards these pioneer cells, but within hours begin to merge in head-to-tail fashion to create large streams of cells moving together towards what has now become a cAMP-emitting aggregation center (Ishikawa-Ankerhold and Müller-Taubenberger, 2019; Nichols et al., 2015). Figure 6, column 1, shows a representative “streaming assay” for WT cells, where large streams had formed by ~6 hours, and aggregation was approaching completion by ~14 hours (see also Movie 3). In sharp contrast, CARMIL-GAP null cell line M1 failed to form streams, making only small cell aggregates by ~17 hours (Figure 6, column 2; see also Movie 4) (a similar result was seen with KO line M2; data not shown). Importantly, the re-expression of CARMIL-GAP in M1 cells largely rescued this defect in streaming, as large streams were apparent by 8 hours (Figure 6, column 3; see also Movie 5 and Figure S5, Panels A1-A4, which shows that GFP-CARMIL-GAP localizes to the leading edge of complemented, ripple-stage null cells). Of note, a Western blot of ripple-stage cells showed that null cells exhibit approximately normal levels of the cAMP receptor CAR1 (Figure S6), arguing that their defect in streaming is most likely not due to an inability to sense cAMP (although see Discussion).

**Figure 6.**
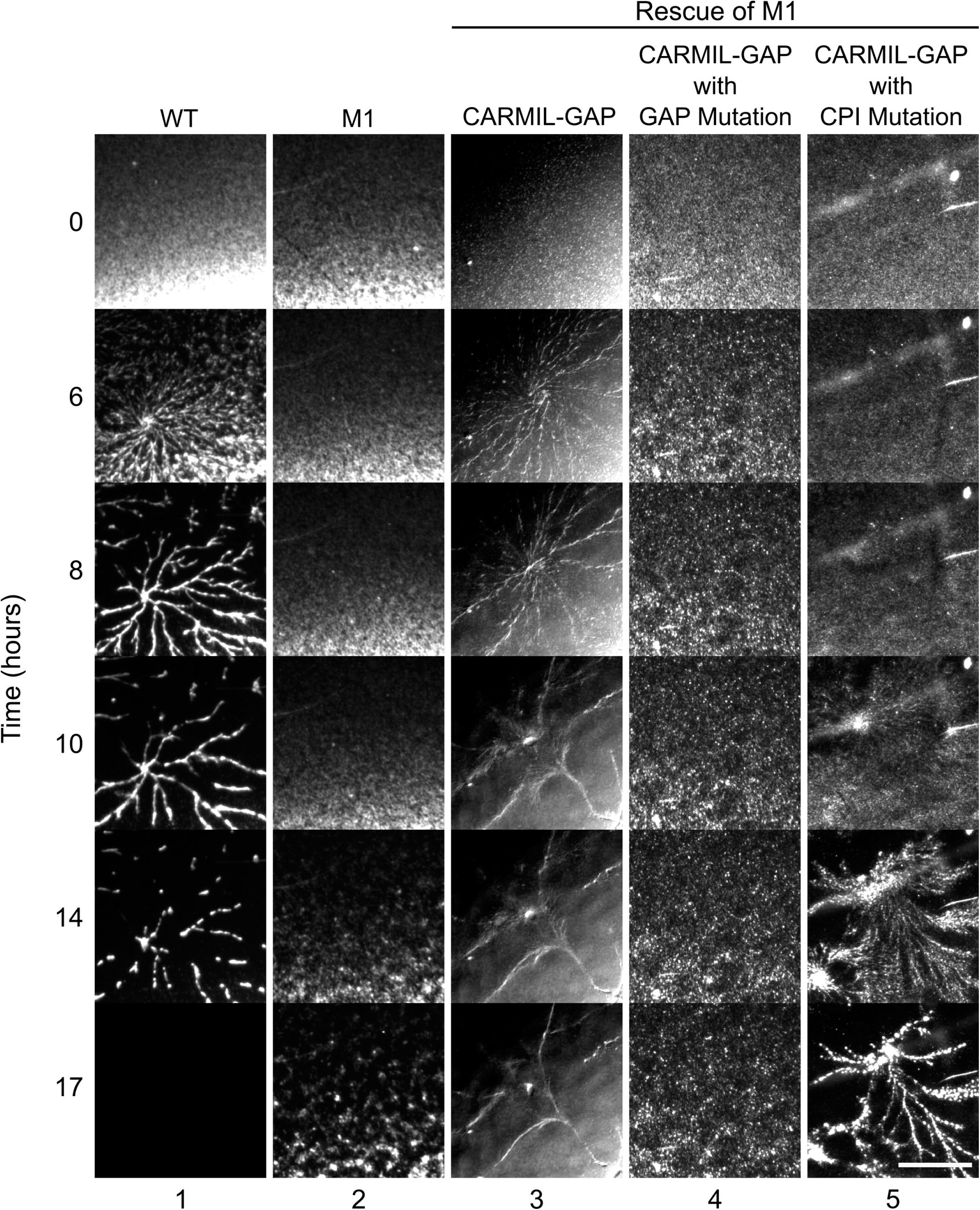
CARMIL-GAP null cells exhibit a defect in chemotactic streaming. Shown are images at 0, 6, 8, 10, 14 and 17 hours of streaming assays for WT and M1 KO cells, and for M1 KO cells that were rescued with GFP-tagged versions of CARMIL-GAP, CARMIL-GAP containing the GAP domain mutation, or CARMIL-GAP containing the CPI domain mutation. While the images are from a single experiment, they are representative of three independent experiments. Mag bar: 1 mm.

To address the underlying cause of the defect in chemotactic aggregation, we starved WT cells and KOs M1 and M2 at high density on black filters until “ripple stage”, which took ~5 hours for both WT and KO cells. At this stage, *Dictyostelium* amoebae exhibit their highest rate of motility, which is about two to four times faster than the rate exhibited by vegetative cells (Hoeller and Kay, 2007; Jung et al., 2016; Jung et al., 1996; Varnum et al., 1986). Ripple-stage cells were harvested by trituration, allowed to attach at low density on chamber slides, and the centroid of every cell in the field of view determined every 15 seconds for 15 minutes to obtain motility rates. Representative path plots for WT cells and CARMIL-GAP null cell lines M1 and M2 (Figure 7, Panels A1-A3, respectively) suggested that CARMIL-GAP null cells were significantly slower (see also Movies 6, 7 and 8). Indeed, quantitation showed that the average rate of motility for CARMIL-GAP null cells was 45.2% (M1) and 44.1% (M2) that of WT cells (Figure 7, Panel B; see also Table 1).

**Figure 7.**
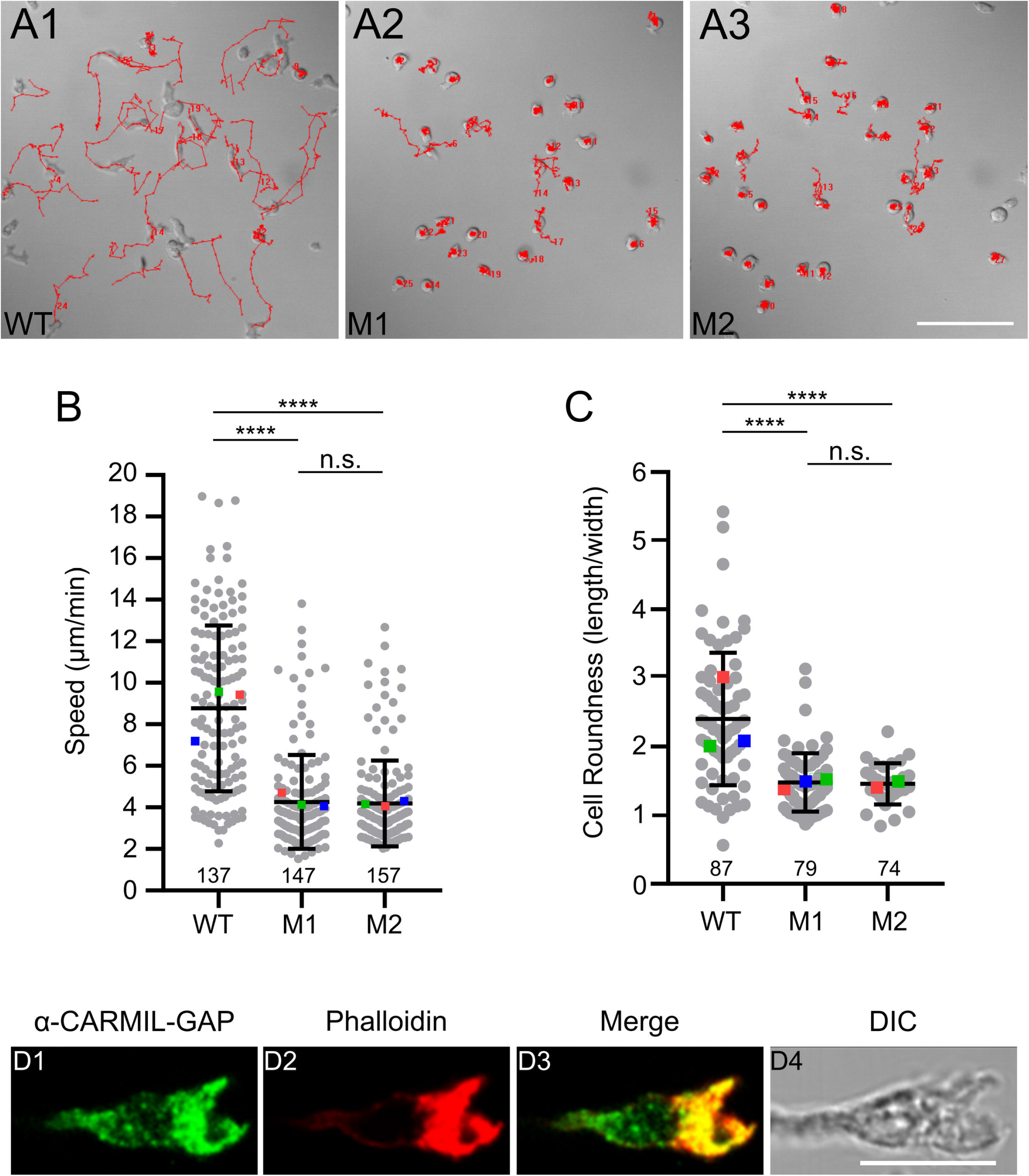
Ripple-stage CARMIL-GAP null cells exhibit defects in cell motility and polarization. (Al-A3) Shown are representative path plots of the random motility exhibited over 15 minutes by ripple-stage WT (Al), M1 KO (A2), and M2 KO (A3) cells. (B) Speeds of ripple-stage WT, M1 KO, and M2 KO cells (see also Table 1). The number of cells scored from three independent experiments is shown at the bottom of each bar, and the mean values for the three experiments performed are indicated by the red, green and blue squares. (C) Cell roundness values (length/width, with 1.0 being perfectly round) exhibited by ripple-stage WT, M1 KO, and M2 KO cells (see also Table 1). The number of cells scored over three independent experiments is shown at the bottom of each bar, and the mean values for the three experiments performed are indicated by the red, green and blue squares. (D1-D4) Image of a representative *Dictyostelium* cell undergoing chemotaxis (to the right) and stained for endogenous CARMIL-GAP (Dl) and F-actin (02) (03 and 04 show the merged and DIC images, respectively). Mag bars: 100 μm in Panel A3 and 10 μm in Panel 04.

During chemotactic aggregation, the fast speed of amoeba is associated with an elongated, highly-polarized shape that aligns with the direction of migration. Higher magnification images of individual cells in the streaming assays performed in Figure 5 showed that CARMIL-GAP null cells appeared on average to be much less polarized than control cells. To quantitate this, we measured the ratio of cell length to cell width using still images from the movies used to determine the motility rates of ripple stage cells. Figure 7, Panel C, shows that both KO M1 and KO M2 are indeed significantly less polarized than WT cells (see also Table 1). Finally, we found that endogenous CARMIL-GAP concentrates along with F-actin in the leading edge pseudopods of chemotaxing cells, as expected (Figure 7, Panels D1-D4).

### While the CPI and GAP domains of CARMIL-GAP both contribute to its function, the GAP domain plays a more significant role

To define the relative contributions that CARMIL-GAP’s CPI domain (CP regulation) and GAP domain (Rac1a regulation) make to its overall function, we complemented CARMIL-GAP null cell line M1 with two mutant versions of CARMIL-GAP expressed as GFP fusions. In one version referred to as “CARMIL GAP with GAP Mutation”, the invariant arginine residue that is required for robust GAP activity was changed to a function-blocking alanine residue. In the other version referred to as “CARMIL-GAP with CPI Mutation”, the invariant arginine residue that is essential for CP binding was changed to a function-blocking glutamate residue. With regard to the phagocytosis of bacteria, Figure 4, Panel C, shows that the GAP domain mutant was completely incapable of rescuing CARMIL-GAP null cell line M1, while the CPI domain mutant partially rescued these cells (the value obtained was in between, and significantly different from, the values for both WT cells and null cell line M1; see also Table 1). With regard to chemotactic streaming, column 4 in Figure 6 shows that the GAP domain mutant appeared completely incapable of rescuing CARMIL-GAP null cell line M1 (see also Movie 9), while column 5 in Figure 6 shows that the CPI domain mutant partially rescued these cells (they did make large streams but took much longer (~14 hours) to do so) (see also Movie 10). Importantly, both mutant GFP-CARMIL-GAP proteins localized to phagocytic cups (Figure S4, Panels B1-B4 and C1-C4) and the leading edge of crawling, ripple-stage cells (Figure S5, Panels B1-B4 and C1-C4). Moreover, both mutant proteins were expressed in complemented null cells at levels exceeding that of wild type GFP-CARMIL-GAP in complemented null cells (which exhibit complete rescue) (Figure S3; compare the GFP-CARMIL-GAP signal in lanes 4 and 5 to lane 3). These results argue that the inability of the GAP mutant to rescue either phagocytosis or streaming, and the inability of the CPI mutant to fully rescue these behaviors, is due to their functional defects rather than to miss localization or insufficient expression. We conclude, therefore, that while both domains contribute significantly to CARMIL-GAP function, the GAP domain plays the more significant role. This conclusion adds to the growing evidence that CARMIL proteins regulate actin dynamics by regulating signaling pathways as well as CP, and that the continued cycling of Rho GTPases between their GTP and GDP bound states through the coordinated action of their GEFs and GAPs and is often required to drive Rho-dependent biological processes forward.

## DICUSSION

CARMIL proteins serve as important regulators of actin-dependent cellular processes by virtue of their ability to regulate Capping Protein (Edwards et al., 2014; Mullins et al., 2018). Evidence is accumulating, however, that they also regulate these processes by interacting with signaling molecules (Stark et al., 2017). Here we showed that *Dictyostelium* CARMIL-GAP is responsible for such dual regulation but as a single protein. Moreover, we determined the functional significance of these two activities by complementing CARMIL-GAP null cells with versions of CARMIL-GAP that lack either GAP activity toward Rac1a or the ability to regulate CP. While both activities were found to contribute significantly to CARMIL GAP’s ability to support phagocytosis and chemotactic streaming, its GAP activity towards Rac1a was the more important of the two. Specifically, CARMIL-GAP lacking GAP activity was completely incapable of rescuing the defect in phagocytosis and yielded only a very modest rescue of the defect in chemotactic streaming, while CARMIL-GAP lacking the ability to regulate CP rescued both processes to a significant extent, although not completely. For this particular CARMIL, therefore, its ability to regulate a signaling pathway contributes more to its overall cellular function than its ability to regulate CP. While this conclusion highlights the functional significance of CARMIL-dependent effects on signaling pathways, the relatively modest role played by CARMIL-GAP’s CPI domain should be considered in the context of possible functional redundancy, as CARMIL-GAP null cells still contain CARMIL and at least one other protein containing a CPI domain (see the legend to Figure S1 for details). It is also important to note that while the discussion below focuses on the role of CARMIL-GAP’s GAP domain in regulating Rac1a, we cannot exclude the possibility that this domain regulates additional Rho-related GTPases (e.g. Rac1E), and that their miss regulation contributes to the defects in actin-dependent processes exhibited by CARMIL-GAP null cells. In a similar vein, we cannot exclude the possibility that the profound defect in streaming exhibited by cells lacking CARMIL-GAP’s GAP activity is due at least in part to the miss regulation of GTPases required for progression of *Dictyostelium’s* developmental program. The pronounced defect in phagocytosis exhibited by null cells cannot be attributed, however, to defects in this developmental program.

Our study focused on Rac1a as the target of CARMIL-GAP’s GAP activity, as 2 of the 6 Rac1-related peptides we obtained were specific to this isoform, and none where specific to Rab1b or Rac1c. That said, Rac1b and Rac1c could also be targets given that 4 of the 6 peptides in our mass spec data are also present in these two isoforms. Any GAP activity towards Rac1b and Rac1c would probably not be that consequential, however, as Rac1a is expressed at vastly higher levels than Rac1b and Rac1c in both vegetative and starved cells (Fey et al., 2013).

We assume that CARMIL-GAP’s GAP domain functions together with one or more GEF partner(s) to regulate the nucleotide state of Rac1a in such a way as to promote phagocytosis and chemotactic streaming *(Dictyostelium* contains 46 genes encoding conventional RhoGEFs; (Rivero and Xiong, 2016)). With regard to the protein or proteins downstream of Rac1a whose activity is regulated by CARMIL-GAP’s GAP activity, one major candidate is the pentameric WAVE Regulatory Complex (WRC), which triggers the formation of branched actin networks by the Arp2/3 complex by coupling active Rac in the plasma membrane to the activation of SCAR/WAVE, a nucleation promoting factor (NPF) for the Arp2/3 complex (Davidson and Insall, 2011; Davidson and Insall, 2013; Mullins et al., 2018; Pollitt and lnsall, 2009; Rivero and Xiong, 2016; Rotty et al., 2013; Schaks et al., 2019; Swaney and Li, 2016). This pathway likely underlies the effect that CARMIL-GAP’s GAP activity has on the process of chemotactic streaming, as the leading-edge pseudopods driving streaming in *Dictyostelium* are known to be created by Arp2/3 complex-dependent branched actin nucleation downstream of Rac1, the WRC and the NPF SCAR/WAVE (Davidson et al., 2018; Rivero and Xiong, 2016; Schaks et al., 2018; Veltman et al., 2012). This pathway may also underlie the effect that CARMIL-GAP’s GAP activity has on the process of phagocytosis, since Rac1 also contributes to the formation of the branched actin networks that comprise much of the phagocytic cup (in this case through the NPF WASP (Davidson et al., 2018; Jeon and Jeon, 2020) as well as the NPF SCAR/WAVE (Seastone et al., 2001). Finally, CARMIL-GAP’s GAP activity may contribute to phagocytosis and chemotactic streaming by regulating additional down-stream effectors of Rac1 (e.g. PAK kinases, formins, IQGAP; (Buckley et al., 2016; Rivero and Xiong, 2016).

An obvious question is why the deletion of CARMIL-GAP does not actually promote phagocytosis and chemotactic streaming given that GAPs push Rho-related GTPases into their GDP-bound “off” state. Indeed, past studies have raised similar questions regarding the regulation of Rho-dependent cellular processes by GEF/GAP pairs based on the simplified view that the GEF should promote the process by increasing the amount GTP-bound Rho, while the GAP should inhibit the process by decreasing the amount of GTP-bound Rho. An early “crack” in this simplified view came from studies showing that dominant active versions of Rho-related GTPases often cannot support biological processes (reviewed in (Parrini and Camonis, 2011)). This crack has continued to widen, as numerous mechanistic studies have revealed a variety of ways in which GEF and GAP activities are both employed to promote Rho-dependent cellular processes (reviewed in (Denk-Lobnig and Martin, 2019)). For example, instances exist where both activities act at the same time in different places, at different times in the same place, and at the same time and in the same place. Importantly, these variations yield variations in the temporal and/or spatial control of the Rho GTPase’s nucleotide state that are required by specific biological processes. For example, oscillations in the nucleotide state of RhoA created by the sequential actions of a RhoA GEF and a RhoA GAP drive the pulsatile contractions of medioapical actomyosin networks that are required for the apical constriction of epithelial cells (Mason et al., 2016). The one constant in all of these studies is that GEFs and GAPs are both required in some fashion or another for the proper execution of Rho-dependent biological processes.

One example of GEF/GAP coordination that is particularly relevant to our study was recently provided by Schlam and colleagues (Schlam et al., 2015), who showed that the phagocytosis of large, IgG-coated particles by macrophages requires the activity of several GAPs for Rac1 and Cdc42, whose activation by GEFs is also required for Fc receptor-mediated phagocytosis. In this case, the GAPs appeared to be driving actin network disassembly at the base of the phagocytic cup even as the cup was still undergoing GEF-dependent pseudopod extension around the particle. The authors suggested that this localized, GAP-dependent actin network disassembly serves to promote phagocytosis by recycling limiting components to the growing tips of advancing pseudopods, as well as by clearing a path for final particle internalization. In many ways, then, this is an example of “same time, different place” GEF/GAP coordination. The defect in the ability of CARMIL-GAP null cells to phagocytose a large particle (yeast), where events commonly reversed part way through the engulfment process, seems in line with the results of Schlam et al, although CARMIL-GAP is not restricted to the base of the phagocytic cup like the GAPs imaged in that study.

Another common form of GEF/GAP coordination is having them act at the same time and in the same place. While this might seem at first blush to create a futile cycle, what it actually creates is a dynamic cycling of the Rho’s nucleotide state between its active and inactive forms (Denk-Lobnig and Martin, 2019). Importantly, such dynamic cycling can be employed by the cell to tune Rho signaling to an optimal level, as well as to rapidly adjust this level to meet varying functional demands. The RhoA-dependent regulation of the actomyosin flows that establish left: right asymmetry in C. *elegans* embryo (Schonegg et al., 2007; Schonegg and Hyman, 2006), and the Rac1-dependent regulation of the branched actin networks that define dendritic spine morphology (Um et al., 2014), represent examples where Rho-related GTPases are controlled by GEFS and GAPs that co-localize and function simultaneously. By analogy, CARMIL-GAP and its partner GEF may function at the same time and in the same place to control the Rac1a-dependent assembly and subsequent turnover of the branched actin networks that comprise the phagocytic cup and the leading edge of migrating cells.

Assuming CARMIL-GAP and its partner GEF do function at the same time and in the same place to drive the rapid cycling of Rac1a’s nucleotide state, how might this serve to promote particle engulfment and cell migration? In thinking about this question, it is important to consider the accumulating evidence that CARMIL proteins shuttle between two states: a folded, inactive form in the cytoplasm that cannot regulate CP, and an unfolded, active form at the plasma membrane that can regulate CP (Fujiwara et al., 2014; Uruno et al., 2006). While the molecules that recruit CARMIL proteins to the plasma membrane have not been identified with absolute certainty, one likely candidate for vertebrate CARMILs is active Rac1, which would interact with these CARMILs via their CRIB domain-like sequences (Fujiwara et al., 2014). Importantly, the recruitment of CARMILs to the plasma membrane by active Rac (and possibly other molecules like polyphosphoinositides) is thought to trigger their unfolding and activation, thereby allowing them to begin converting sequestered CP (CP bound to V-1) into CARMIL: CP complexes that then cap nascent branched actin filaments (Fujiwara et al., 2014; Mullins et al., 2018). If CARMIL-GAP is also recruited to the plasma membrane by active Rac1a, then its GAP activity towards Rac1a could drive repeated rounds of CARMIL-GAP recruitment to, and release from, the plasma membrane when combined with the simultaneous activity of a Rac1a GEF. Given the likelihood that membrane-bound CARMIL-GAP also collaborates with V-1 in *Dictyostelium* (Jung et al., 2016) to drive the capping of nascent filament barbed ends required for branched actin network formation, then the rapid cycling of CARMIL-GAP on and off the plasma membrane might serve to promote the advance of pseudopods during phagocytic particle engulfment and leading edge extension.

## MATERIALS AND METHODS

### Cell biological methods

*Dictyostelium* strain AX3 was grown in HL5 medium, transformed by electroporation, and stable clonal transformants were isolated as described previously (Jung et al., 1996). The growth of *Dictyostelium* on plates containing *Klebsiella aerogens* bacteria (plaque assay) was performed as described previously (Jung et al., 1996). The initial rate of uptake of 1 μm FITC-latex beads by phagocytosis, and the rate of macropinocytosis of FITC-dextran, were performed as described previously (Jung et al., 2001). Streaming assays, measurements of the speed and polarity of ripple-stage cells, measurements of growth rates and the number of nuclei/cell, and measurements of cellular F-actin content by FACs analysis of cells stained with FITC-phalloidin, were all performed as described previously (Jung et al., 1996). The preparation of whole cell extracts, SDS-PAGE, and Western blotting were performed as described previously (Jung et al., 2001). Fixing, immunostaining and imaging *Dictyostelium* on a Zeiss LSM 780 microscope equipped with a 63X 1.4-NA objective (or a 40X 1.2 NA for the motility assays) were performed as described previously (Jung et al., 1996).

### Vectors

Wild type CARMIL-GAP, CARMIL-GAP containing the arginine to alanine point mutation at residue 737 in the GAP domain, and CARMIL-GAP containing the arginine to glutamate point mutation at residue 989 in the CPI domain, were cloned with EcoR1 ends using standard techniques, sequenced confirmed, and expressed as GFP fusions using the vector pDEX H (Jung et al., 1996) Transformation and the selection of stable transformants using G418 were performed as described previously (Jung et al., 2016).

### Reagents

The polyclonal antibody to the alpha subunit of *Dictyostelium* Capping Protein was a generous gift of John A. Cooper (Washington University, St. Louis). Blasticidin-S and G418 were purchased from Sigma. Labeled phalloidins and secondary antibodies and the protein molecular weight marker were purchased from Thermo Fisher Scientific. The mouse monoclonal antibody against CAR1 was a generous gift of Carol Parent (National Cancer Institute, Bethesda). Protein concentrations were determined by Bradford assay (Bio-Rad).

### CARMIL-GAP antibody

A peptide corresponding to CARMIL-GAP residues 939 to 953 was synthesized, conjugated with KLH, and injected into rabbits using standard techniques (Wang et al., 2010). Rabbit sera obtained after primary immunization and three boosts was purified by absorption against CARMIL-GAP null cell extracts as previously described (Jung et al., 1996).

### CARMIL-GAP knockout

A linear gene-disruption fragment designed to promote a double-crossover gene replacement event was created by fusing nucleotides 1080 to 1591 and nucleotides 2193 to 2715 of the *Dictyostelium* CARMIL-GAP genomic sequence (DictyBase Gene ID No. DDG_G0290439) to the 5⍰ and 3⍰ ends of the Blasticidin resistance cassette in plasmid Bsr2 (Sutoh, 1993) respectively, using the same approach as described previously for knocking out *Dictyostelium* MyoJ (Jung et al., 2009). The introduction of this linear fragment into AX3 cells by electroporation, and the isolation of Blasticidin-resistant clones by serial dilution in 96-well plates, were performed as described previously (Jung et al., 2009) CARMIL-GAP null cell lines M1 and M2 were identified by Western blotting using the CARMIL-GAP antibody.

### GAP domain pulldown

The GAP domain of CARMIL-GAP (residues 715-858) was synthesized by Blue Heron Inc. as an EcoR1/Xho1 fragment using *E. coli* codon bias and a QSGAG spacer between GST and the protein, and then cloned into pGST-Tev Parallel #2 using standard techniques to create GST-GAP. The expression of the GST-GAP fusion protein in *E.coli* strain BL-21-RILP (Stratagene) and its purification using Glutathione Sepharose 4B were performed as described previously (Jung et al., 2001; Jung et al., 2016). For the pull down, Glutathione Sepharose 4B beads loaded with GST-GAP were incubated at 4°C for 2 hours with a *Dictyostelium* whole cell extract that had been dialyzed into 1X TBS containing 5 μM GTPyS. After five washes with 1X TBS, bound proteins were eluted with high salt buffer (5X TBS), concentrated using a Ultracel-10 (10 kDa) Amicon filter, and subjected to mass spec analysis.

### Liquid chromatography (LC) tandem mass spec (MS) analysis

Protein identification by LC-MS/MS analysis of peptides was performed using an Orbitrap Fusion Lumos Tribid mass spectrometer (Thermo Fisher Scientific, San Jose, CA) interfaced with an Ultimate 3000 Nano-HPLC apparatus (Thermo Fisher Scientific, San Jose, CA). Peptides were fractionated by EASY-Spray PepMAP RPLC C18 column (2μm, 100A, 75 μm x 50cm) using a 120-min linear gradient of 5-35% acetonitrile in 0.1% formic acid at a flow rate of 300 nl/min. The instrument was operated in data-dependent acquisition mode using Fourier transform mass analyzer for one survey MS scan on selecting precursor ions followed by 3 second data-dependent HCD-MS/MS scans for precursor peptides with 2-7 charged ions above a threshold ion count of 10,000 with normalized collision energy of 37%. Survey scans of peptide precursors from 300 to 2000 m/z were performed at 120k resolution and MS/MS scans were acquired at 50,000 resolution with a mass range m/z of 100-2000.

### Protein identification and data analysis

All MS and MS/MS raw spectra from each set were processed and searched using Mascot algorithm within the Proteome Discoverer 1.4 (PD 1.4 software, Thermo Scientific). Precursor mass tolerance was set at 20 ppm and fragment ion mass tolerance was set at 0.05 Da. Trypsin was selected as the enzyme, with two missed cleavages allowed. Carbamidomethylation of cysteine was used as fixed modification. Deamidation of glutamine, deamidation of asparagine and oxidation of methionine were used as variable modifications. The *Dictyostelium discoedium* sequence database from Swiss Sprot was used for the database search. Identified peptides were filtered for maximum 1% FDR using the Percolator algorithm in PD 1.4 along with additional peptide confidence set to medium. The final lists of protein identification/quantitation were filtered by PD 1.4 with at least 2 unique peptides per protein identified with medium confidence. For the quantitation, a label-free approach was used, where the area under the curve for the precursor ions is used to calculate the relative fold change between different peptide ions.

### CPI domain pulldown

The CPI domain of CARMIL-GAP (residues 965 to 1005) was synthesized by Blue Heron Inc. as an EcoR1/Xho1 fragment using E. coli codon bias and a QSGAG spacer between GST and the protein, and then cloned into pGST-Tev Parallel #2 using standard techniques to create GST-CPI. A second version in which the essential arginine at position 989 was changed to a glutamate (GST-CPI-R⍰E) was also synthesized. The expression of the GST-CPI and GST-CPI-R⍰E fusion proteins in E. coli strain BL-21-RILP and their purification using Glutathione Sepharose 4B were performed as described previously (Jung et al., 2001). The pulldown of CP using these two fusion proteins was performed exactly as described previously for the pulldown of CP by GST-V1 (Jung et al., 2016).

### GAP assay

*Dictyostelium* Rac1a was synthesized by Blue Heron Inc. as an EcoRl/Xhol fragment using E. coli codon bias and a QSGAG spacer between GST and the protein, and then cloned into pGST-Tev Parallel #2 using standard techniques to create GST-Rac1a. The expression of the GST-Rac1a in E. *coli* strain BL-21-RILP its purification using Glutathione Sepharose 4B were performed as described previously (Jung et al., 2016; Jung et al., 2001) GAP assays were performed as described previously (Faix et al., 1998) except that GTP hydrolysis was quantitated using CytoPhos reagent (Cytoskeleton Inc) to measure the amount of free phosphate. Briefly, purified GST-Rac1a was converted into its GTP-bound form by incubation with a 50-fold molar excess of GTP in the presence of 40 mM EDTA and 200 mM (NH4)2504 at 4°C for 1 hour, desalted into 20 mM Tris-HCl (pH 7.5), 5 mM MgCl2, and 1 mM DTT to remove free nucleotide, and concentrated by Amicon filtration. The amount of phosphate released by GTP hydrolysis after a 10 minute incubation at 20°C was determined for Rac1a alone, Rac1a plus GST-GAP, and Rac1a plus GST-GAP-RΔA using CytoPhos reagent according to the manufacturer’s instructions.

### Bacteria phagocytosis assays

The FACS-based phagocytosis assay employing pHRodo Red-labeled *Klebsiella aerogenes* (which fluoresce brightly when subjected to the low pH inside acidified phagolysosomes) was performed exactly as described previously (Pan et al., 2018a; Pan et al., 2016). Briefly, *Klebsiella aerogenes* labeled with pHrodo Red dye (Life Technologies) were incubated at 22°C and 150 rpm with WT and CARMIL-GAP KO cells suspended in phosphate buffer (7.4 mM NaH_2_P0 4·H_2_0, 4 mM Na_2_HP0 _4_·7H_2_0, 2 mM M gCl_2_, 0.2 mM CaCl_2_, pH 6.5) at a ratio of ~100 bacteria per *Dicytostelium* cell. At the indicated times, *Dictyostelium* cells were pelleted by centrifugation, resuspended in an alkaline buffer (50 mM Tris (pH 8.8), 150 mM NaCl) to quench any possible fluorescence coming from non-engulfed bacteria, and the fluorescence signal for pHrodo Red inside *Dictyostelium* determined by flow cytometry using a FACSort flow cytometer (BD Bioscience), Cell Quest software (v. 3.3) and the FlowJo analysis program (v. 10.0.8; Tree Star). The imaging-based phagocytosis assay employing pHRodo Red-labeled *Klebsiella aerogenes* was also performed exactly as described previously (Pan et al., 2016; Pan et al., 2018a). Briefly, *Dictyostelium* cells were allowed to attach to chamber slides (Lab-Tek) and then incubated with pHrodo Red-labeled *bacteria* in phosphate buffer at a ratio of ~1 *Dictyostelium* to 50 bacteria. After 15 minutes, the bacteria-containing buffer in the chamber slide was replaced with the alkaline buffer described above to halt further bacteria engulfment and quench any fluorescence coming from un-engulfed bacteria. The number of engulfed bacteria per *Dictyostelium* cell was then determined by imaging using a Zeiss LSM 880 confocal microscope equipped with 60X 1.3 NA Plan-Neofluar objective lens.

### Yeast particle phagocytosis

Heat-killed S. cerevisiae (Invivogen) were labeled with Tetramethyl Rhodamine Isothiocyanate (TRITC) as described previously (Rivero and Xiong, 2016) *Dictyostelium* cells were mixed 1: 5 with TRICT-labeled yeast particles in HL5 media and placed in a chambered cover glass. After 10 min at 20°C, the capture of DIC and fluorescence images was commenced at 5 second intervals for 25 to 35 minutes using a Zeiss LSM 780 equipped with a 40 X, 1.2 NA objective. For each event scored, time zero corresponded to contact between a *Dictyostelium* cell and a yeast particle. Failed phagocytic events, which represented the bulk of events scored, corresponded to the subsequent separation of the yeast particle from the *Dictyolstelium* cell. Successful phagocytic events were defined by two criteria: (1) the *Dictyolstelium* cell and the yeast particle remained together for at least 15 minutes (or roughly five times longer than the average time to failure), and (2) examination of every such events correspond to successful phagocytosis events was obtained by examination of every video frame in the last 5 minutes of the 15 minute period for every events judged successful, which showed that the cell-associated yeast particle remained at all times within the 20 footprint of the cell even as it changed shape or migrated

### Statistics

Statistical significance was determined using unpaired t-test and indicated as follows: * = P<0.05, ** = P<0.01, *** = P<0.001, and **** = P<0.0001.

## ACKNOWLEDGEMENTS

We thank Dr. J. Philip McCoy and Ms. Leigh Samsel (National Heart, Lung, and Blood Institute Flow Cytometry Core), Dr. Marjan Gucek and Ms. Sajni Patel (National Heart, Lung, and Blood Institute Proteomics Core), and Dr. Xufeng Wu (National Heart, Lung, and Blood Institute Light Microscopy Core) for their contributions to this work. We also thank Dr. Robert Insall (University of Glasgow) for advice of *Dictyostelium* Rac1 GTPases. Finally, JAH wishes to thank GJ for the 35 years we worked together, and hopes GJ has a long, healthy and relaxing retirement.

**Figure Sl. Alignment of the sequences of CARMIL and CARMIL-GAP.** Sequences corresponding to the domains shared between CARMIL-GAP and CARMIL, and the GAP domain specific to CARMIL-GAP, are indicated by a colored highlighting that matches the color of the domains in the Figure 1 cartoon. Lines indicate identity, colons indicate highly conservative substitutions, and dots indicate moderately conservative substitutions. Of note, it is unclear whether CARMIL-GAP has a bona fide HD domain based on this alignment. Also of note, searches of the *Dictyostelium* genome for proteins other than CARMIL and CARMIL-GAP that contain a CPI domain identified only one other protein (the *Dictyostelium* homolog of the WASH complex subunit FAM21). Finally, searches of the genome also identified a second CARMIL-GAP gene (XP-003280987) that is similar to CARMIL-GAP (including the GAP domain), although it is missing the CPI domain.

**Figure S2. Vegetative CARMIL-GAP null cells do not exhibit defects in growth rate, macropinocytosis, F-actin content, or cytokinesis.**(A) Western blot of whole cell extracts of vegetative AX3 cells ("Veg") and AX3 cells starved for 2, 4, 6, 8, and 10 hours on black filters, probed with the anti-CARMIL-GAP antibody. (B) Growth curve for WT and M1 KO cells. Consistent with this one example, routine maintenance of these cells side by side did not reveal any obvious difference in growth rate. (C) Rate of macropinocytic uptake of FITC-labeled dextran by WT and M1 KO cells (WT 15 min, 21.7 ± 5.6 AU; WT 30 min, 37.1 ± 1.2 AU; WT 45 min, 63.5 ± 2.8 AU; WT 60 min, 82.2 ± 8.1 AU; WT 75 min, 98.5 ± 16.7 AU; WT 90 min, 100.0 ± 0.0 AU; M1 15 min, 22.3 ± 3.5 AU; M1 30 min, 44.8 ± 13.1 AU; M1 45 min, 72.1 ± 23.2 AU; M1 60 min, 79.6 ± 12.7 AU; M1 75 min, 89.3 ± 16.6 AU; WT 90 min, 97.6 ± 11.3 AU; N=3). (D) Total cellular F-actin content for WT and M1 KO cells, as determined by FACs analysis of cells stained with FITC-phalloidin (Jung et al., 2016). (E) Percent of mid-log WT and M1 KO cells with one (green), two (orange), or 3 or more (red) nuclei per cell. The total number of cells scored on three different days is indicated at the top of each bar.

**Figure S3. Western blots of rescued CARMIL-GAP KO M1.** Shown is a Western blot of whole cell extracts of WT cells (lane 1), CARMIL-GAP KO M1 (lane 2), and CARMIL-GAP KO M1 rescued with GFP-CARMIL-GAP (lane 3), GFP-CARMIL-GAP with the GAP mutation (lane 4), and GFP-CARMIL-GAP with the CPI mutation (lane 5), and probed with antibodies to CARMIL-GAP and actin as a loading control (of note, this single blot was cut into two pieces to probe with the two antibodies). The weak band in lanes 3-5 that corresponds in size to CARMIL-GAP most likely reflects the proteolytic cleavage of GFP from the GFP-CARMIL-GAP fusion proteins. The fact that M1 is fully rescued by very modest expression of GFP-CARMIL-GAP is consistent with our unpublished data on CARMIL, where CARMIL null cells are also fully rescued by very modest expression of GFP-CARMIL. Modest expression of GFP-tagged CARMIL and CARMIL-GAP may be sufficient to support the full rescue their null lines because the addition of GFP appears to favor the unfolded, active conformation of these proteins (Uruno et al 2006, Fujiwara et al, 2014, unpublished results).

**Figure S4. GFP-CARMIL-GAP, GFP-CARMIL-GAP with the GAP mutation, and GFP-CARMIL-GAP with the CPI mutation localize to the phagocytic cup in rescued KO M1.** Shown are representative examples of CARMIL-GAP KO M1 cells expressing GFP-CARMIL-GAP (A1-A4), GFP-CARMIL-GAP with the GAP mutation (B1-B4), and GFP-CARMIL-GAP with the CPI mutation (C1-C4) that were incubated with yeast particles as a phagocytic substrate and then fixed and stained with Phalloidin. The faint green signal on the yeast particles in Panels Al, Bl and Cl is due to bleed through from the extremely intense Phalloidin signal on these particles in the red channel. Mag bars: 1.6 μm.

**Figure S5. GFP-CARMIL-GAP, GFP-CARMIL-GAP with the GAP mutation, and GFP-CARMIL-GAP with the CPI mutation localize to the leading edge in rescued, ripple-stage KO M1.** Shown are representative examples of motile, ripple-stage CARMIL-GAP KO M1 cells expressing GFP-CARMIL-GAP (A1-A4), GFP-CARMIL-GAP with the GAP mutation (B1-B4), and GFP-CARMIL-GAP with the CPI mutation (C1-C4) that were fixed and stained with Phalloidin (the cells are aligned so that the hyaline zone at their leading edge (see the DIC image) is pointing up and slightly to the left). Mag bars: 3.4 μm.

**Figure S6. CAR1 Western blot.** Shown is a Western blot of whole cell extracts of ripple-stage WT cells (lane 1), CARMIL-GAP KO M1 (lane 2), and CARMIL-GAP KO M2 (lane 3) probed with antibodies to the cAMP receptor CAR1 and actin as a loading control (of note, this single blot was cut into two pieces to probe with the two antibodies). The blot is presented in such a way as to allow direct comparison with a similar blot published by Jung et al (2016), which included a vegetative cell extract to show that the CAR1 band at ~50 kDa is present only in starved cells, as expected.

**Table S1. Curated list of proteins that bound to GST-GAP.** List of proteins identified by mass spec in the eluate of the GST-GAP affinity column after removing ribosomal proteins, mitochondrial matrix proteins, proteins related to transcription/translation, and any proteins identified by less than three peptides.

## MOVIE LEGENDS

**Movie 1.** A failed yeast phagocytosis event by a WT cell. Images were taken every 5 sec for 2 min and are played back a 7 fps.

**Movie 2.** A successful yeast phagocytosis event by a M1 KO cell. Images were taken every 5 sec for 2 min and are played back a 7 fps.

**Movie 3.** Steaming of WT cells. Images were taken every 10 min for 17 hrs and are played back a 30 fps.

**Movie 4.** Steaming of M1 KO cells. Images were taken every 10 min for 17 hrs and are played back a 30 fps.

**Movie 5.** Steaming of M1 KO cells complemented with GFP-CARMIL-GAP. Images were taken every 10 min for 17 hrs and are played back a 30 fps.

**Movie 6.** Path plot of the random motility of ripple-stage WT cells. Images were taken every 15 sec for 15 min and are played back at 30 fps.

**Movie 7.** Path plot of the random motility of ripple-stage M1 KO cells. Images were taken every 15 sec for 15 min and are played back at 30 fps.

**Movie 8.** Path plot of the random motility of ripple-stage M2 KO cells. Images were taken every 15 sec for 15 min and are played back at 30 fps.

**Movie 9.** Steaming of M1 KO cells complemented with GFP-CARMIL-GAP with the GAP mutation. Images were taken every 10 min for 17 hrs and are played back a 30 fps.

**Movie 10.** Steaming of M1 KO cells complemented with GFP-CARMIL-GAP with the CPI mutation. Images were taken every 10 min for 17 hrs and are played back a 30 fps.

